## Supplemental Figures for "Neuronal Detection Triggers Systemic Digestive Shutdown in Response to Adverse Food Sources"

### Figure 1—figure supplement 1

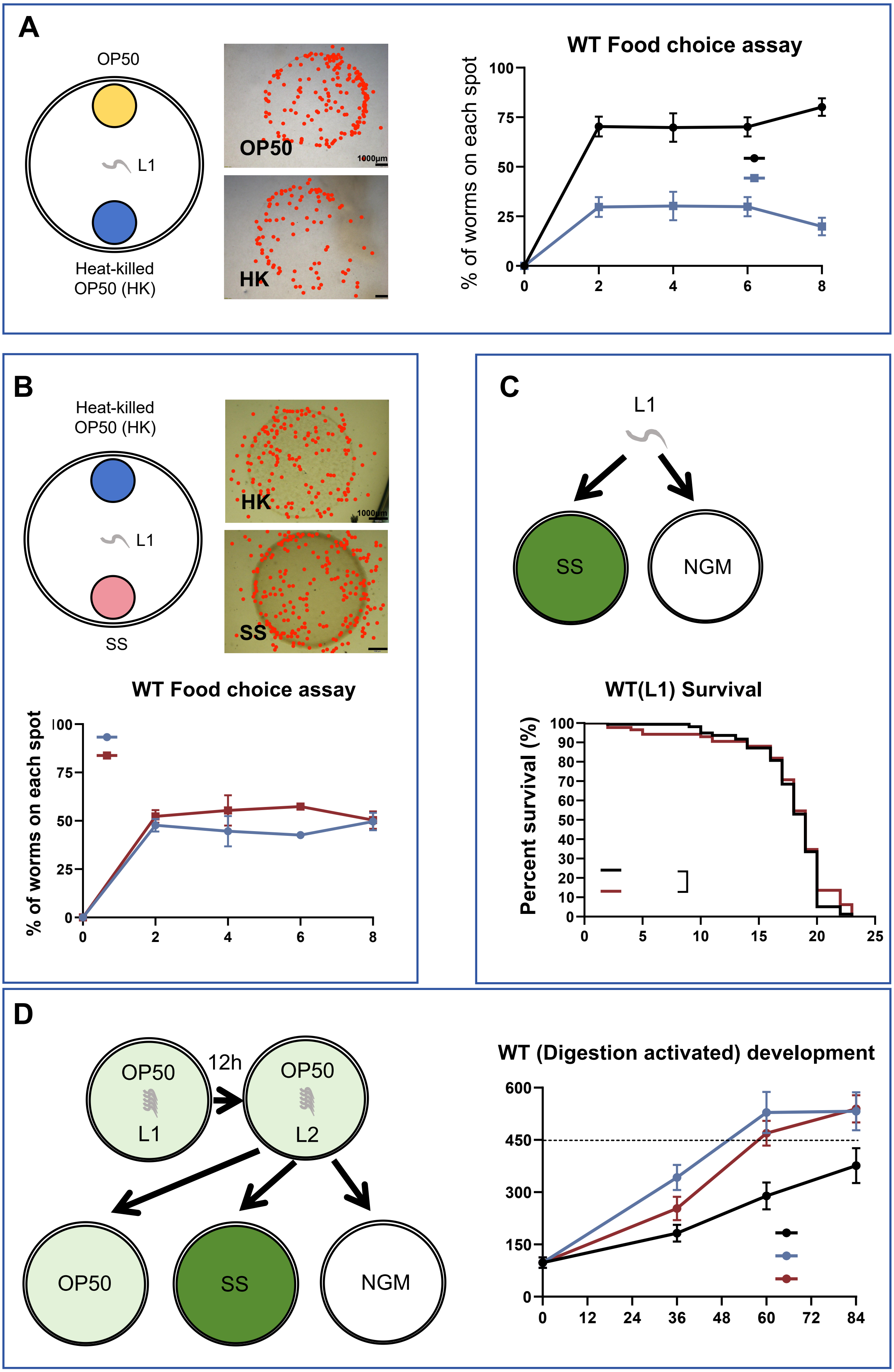

**Figure 1—figure supplement 1. SS is harmful food that animals cannot digest. Related to Figure 1.**

**(A)** Schematic drawing, microscopic images, and quantitative data from the food choice assay. L1 worms were placed at the center spot (origin). OP50 (yellow) and heat-killed OP50 (blue) bacteria were positioned on opposite sides of the plate. The red point indicates the position of each worm. The percentage of worms on each spot was calculated at the indicated times. Data are represented as mean ± SD. Scale bar= 1000 μm. ****$p < 0.0001$; ***$p < 0.001$; **$p < 0.01$ by Student's t-test.

**(B)** Schematic drawing, microscopic images, and quantitative data from the food choice assay. L1 worms were placed at the center spot (origin). Heat-killed OP50 (blue) and SS (red) bacteria were positioned on opposite sides of the plate. The red point indicates the position of each worm. The percentage of worms on each spot was calculated at the indicated times. Data are represented as mean ± SD. Scale bar= 1000 μm. ****$p < 0.0001$; n.s. not significant by Student's t-test.

**(C)** Schematic drawing and quantitative data of the lifespan of animals fed with SS or OP50. L1 worms were seeded on plates with no food (NGM) or SS bacteria to measure lifespan. n.s. not significant by log-rank test.

**(D)** Schematic drawing and quantitative data of the developmental progression of wild-type N2 under different feeding conditions. L1 animals were seeded onto OP50 plates and grown to the L2 stage. L2 animals were then transferred to OP50, SS, or no food (NGM) plates to measure worm length at the indicated time points.

All data are representative of at least three independent experiments.

### Figure 2—figure supplement 1

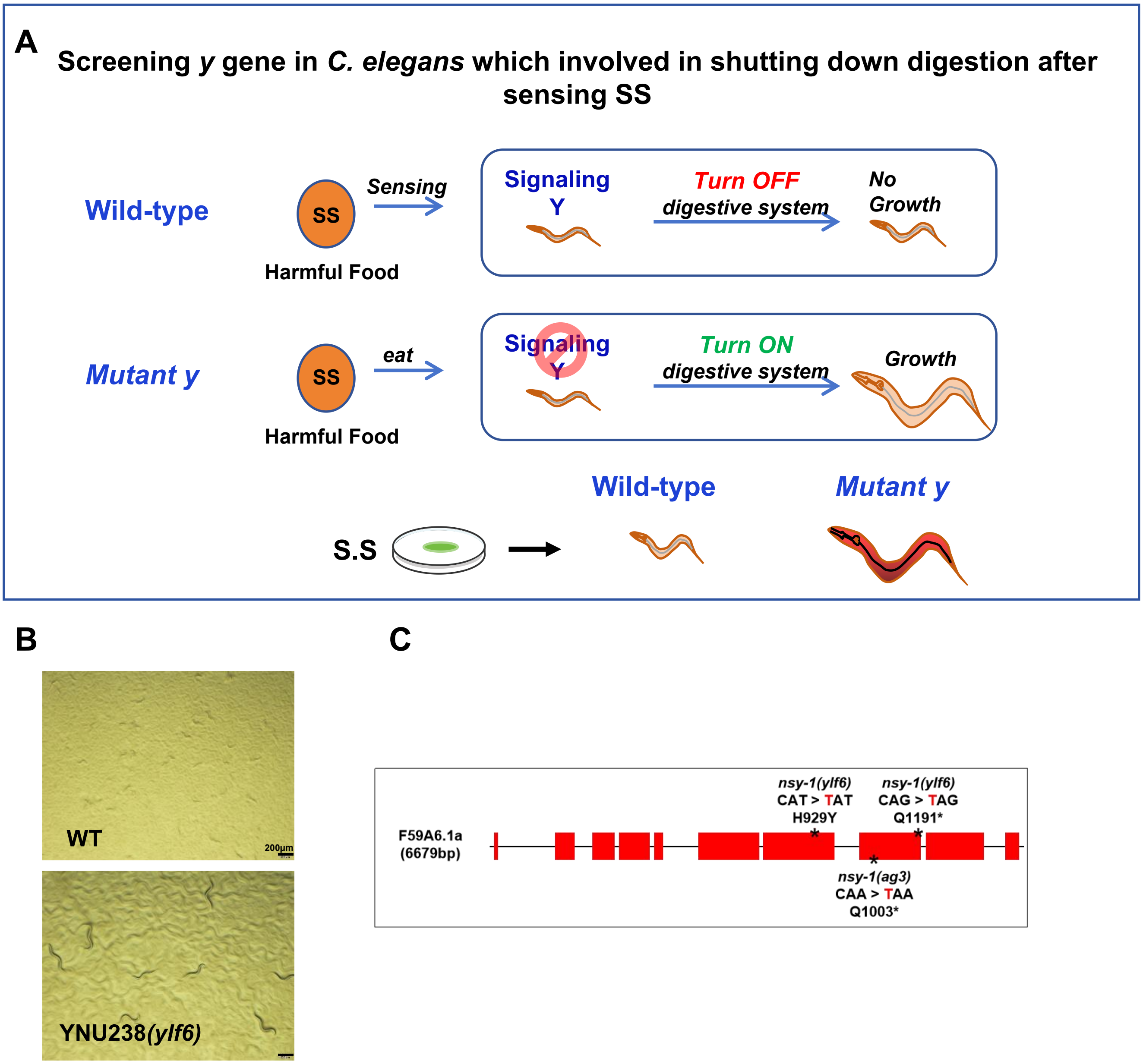


**Figure 2—figure supplement 1. EMS screen to identify genes involved in shutting down digestion of SS.**

**(A)** Schematic illustration of the EMS screen strategy to identify "Y" genes involved in shutting down digestion after sensing SS. In mutants with defects in "Y" genes, digestion of SS is restored, allowing the mutants to grow on SS.

**(B)** Developmental phenotype of wild-type N2 and *ylf6* mutant worms fed with SS bacteria. Scale bar = 200 μm.

**(C)** Schematic drawing showing the mutation sites in *nsy-1(ylf6)* and *nsy-1(ag3)*.

### Figure 3—figure supplement 1

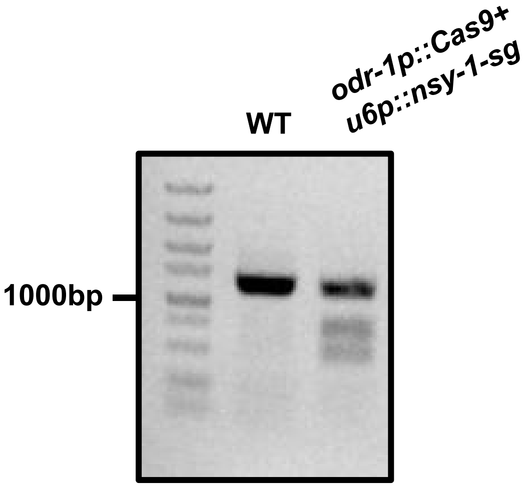


**Figure 3—figure supplement 1. Construction of *nsy-1* specific knockout in AWC neurons using CRISPR-Cas9.**

Representative DNA gels showing NheI digestion of PCR-amplified genomic DNA extracted from wild-type (WT) worms and worms with *nsy-1*-specific knockout in AWC neurons (*odr-1p::Cas9* + *u6p::nsy-1-sg*).

### Figure 3—figure supplement 2

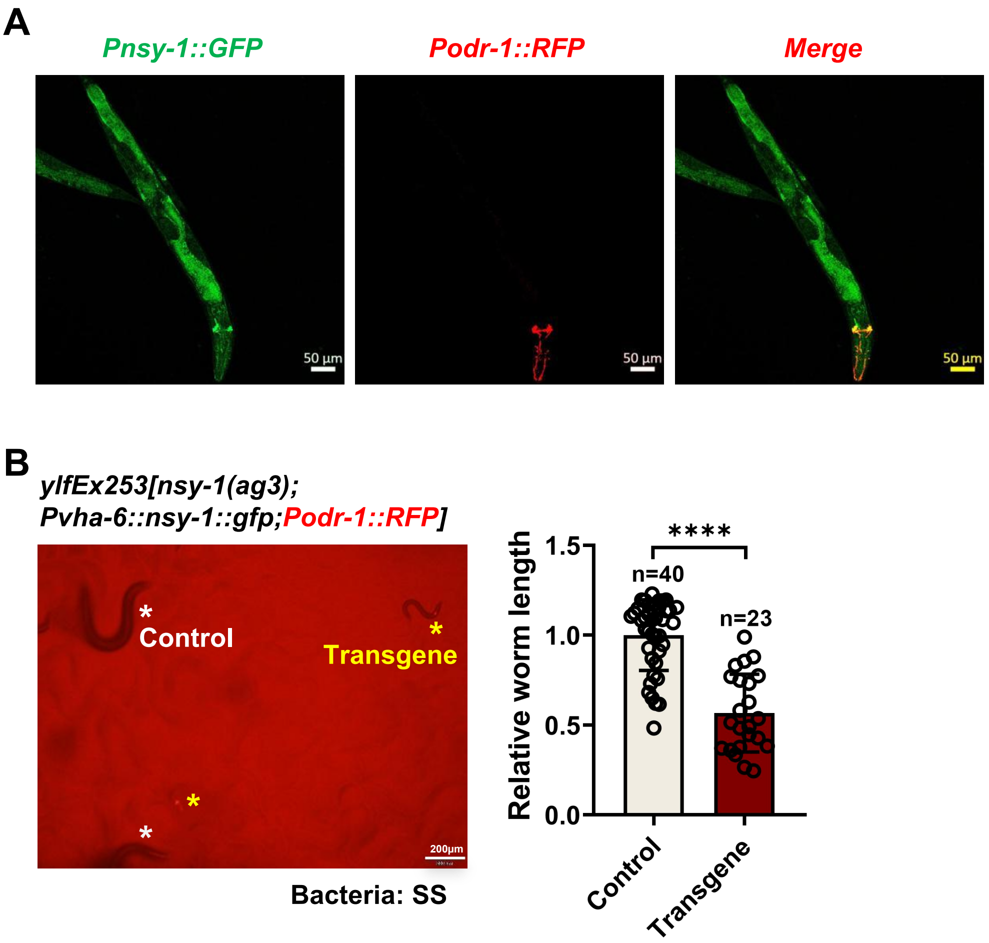


**Figure 3—figure supplement 2. NSY-1 functions in intestine to shut down SS Digestion.**

**(A)** Microscopic image showing the expression pattern of *nsy-1*. *nsy-1* is mainly expressed in the head neurons and intestine. Scale bar = 50 µm.

**(B)** Developmental progression of *nsy-1(ag3)* mutant worms carrying *Pvha-6::nsy-1::gfp* (intestine-specific expression) grown on SS bacteria. Control animals are labeled with white stars, and animals carrying the transgenes (rescued animals) are labeled with yellow stars. Data are represented as mean ± SD. Scale bar = 200 µm. ****p < 0.0001 by Student's t-test. n= number of animals which were scored.

### Figure 3—figure supplement 3

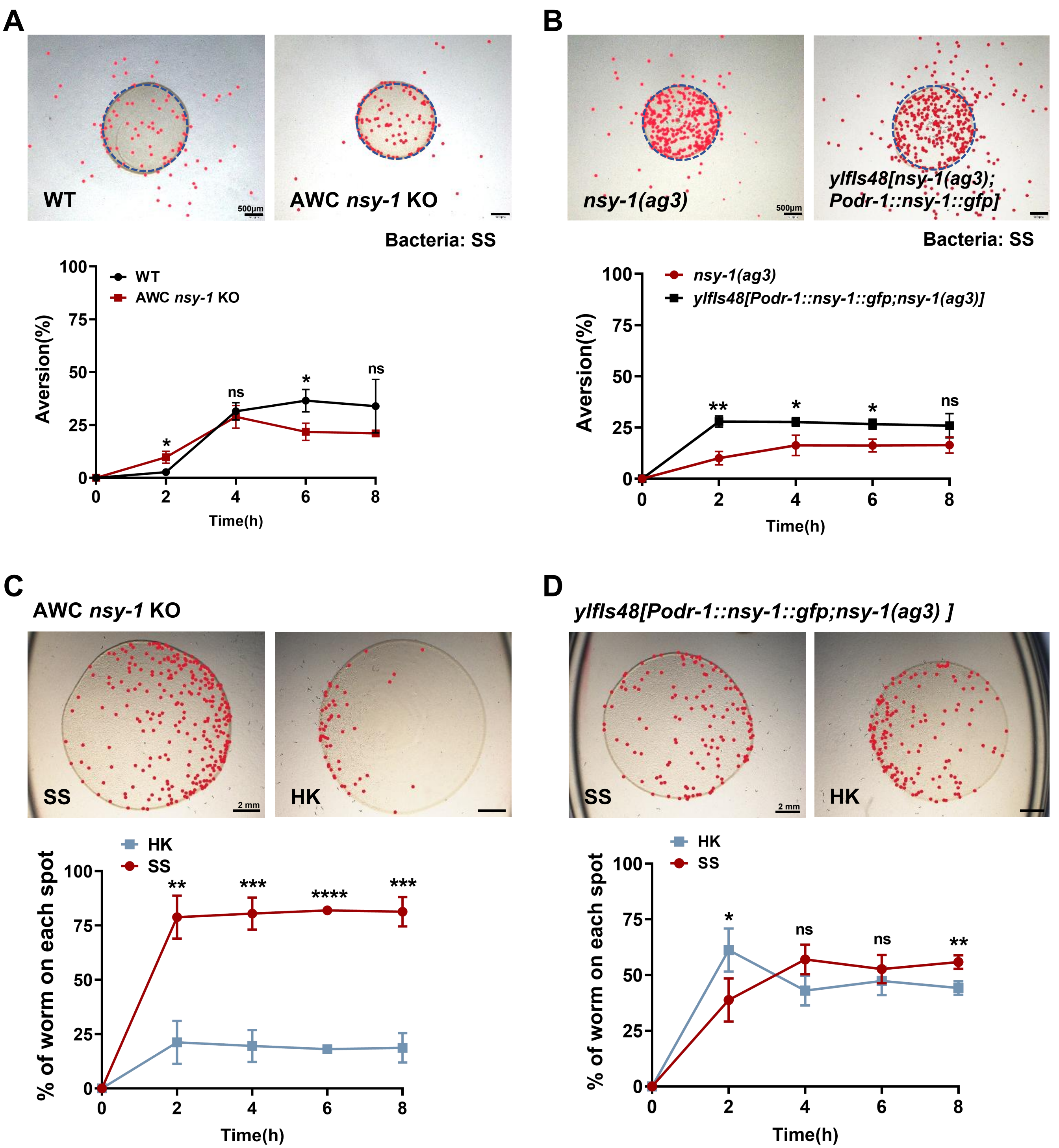

**Figure 3—figure supplement 3. NSY-1 functions in AWC neurons to influences the recognition of SS.**

**(A)** Microscopic images, and quantitative data of the food dwelling/avoidance assay. The animals were scored at the indicated times after L1 worms were placed on the food spot. The blue circle indicates the edge of the bacterial lawn, and the red point indicates the position of each worm. Data are represented as mean ± SD. Scale bar= 500 μm. *p < 0.05 by Student's t-test.

**(B)** Microscopic images, and quantitative data of the food dwelling/avoidance assay. The animals were scored at the indicated times after L1 worms were placed on the food spot. The blue circle indicates the edge of the bacterial lawn, and the red point indicates the position of each worm. Data are represented as mean ± SD. Scale bar = 500 μm. **p < 0.01; *p < 0.05 by Student's t-test.

**(C)** Microscopic images, and quantitative data from the food choice assay. L1 worms were placed at the center spot (origin). Heat-killed OP50 and SS bacteria were positioned on opposite sides of the plate. The red point indicates the position of each worm. The percentage of worms on each spot was calculated at the indicated times. Data are represented as mean ± SD. Scale bar= 2 mm. ****p < 0.0001; ***p < 0.001; **p < 0.01 by Student's t-test.

**(D)** Microscopic images, and quantitative data from the food choice assay. L1 worms were placed at the center spot (origin). Heat-killed OP50 and SS bacteria were positioned on opposite sides of the plate. The red point indicates the position of each worm. The percentage of worms on each spot was calculated at the indicated times. Data are represented as mean ± SD. Scale bar = 2 mm. **p < 0.01; *p < 0.05; n.s. not significant by Student's t-test.

### Figure 3—figure supplement 4

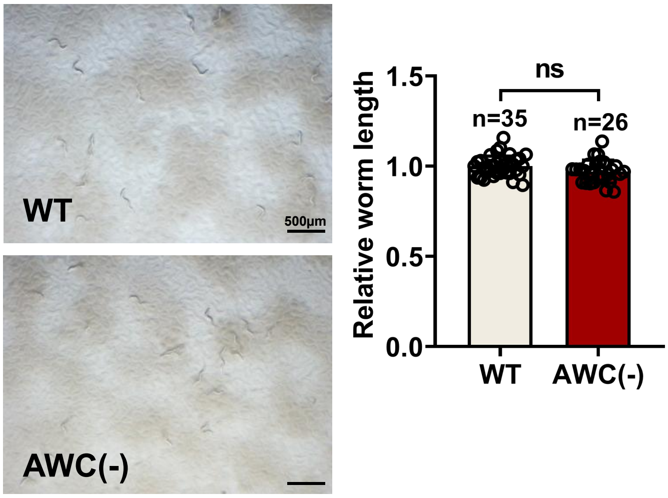


**Figure 3—figure supplement 4. AWC neurons are essential for initiating SS digestion.**

Developmental phenotype of wild-type N2 and AWC(-) worms fed with SS bacteria. Data are represented as mean ± SD. Scale bar = 500 μm. n.s. not significant by Student's t-test. n= number of animals which were scored.

### Figure 4—figure supplement 1

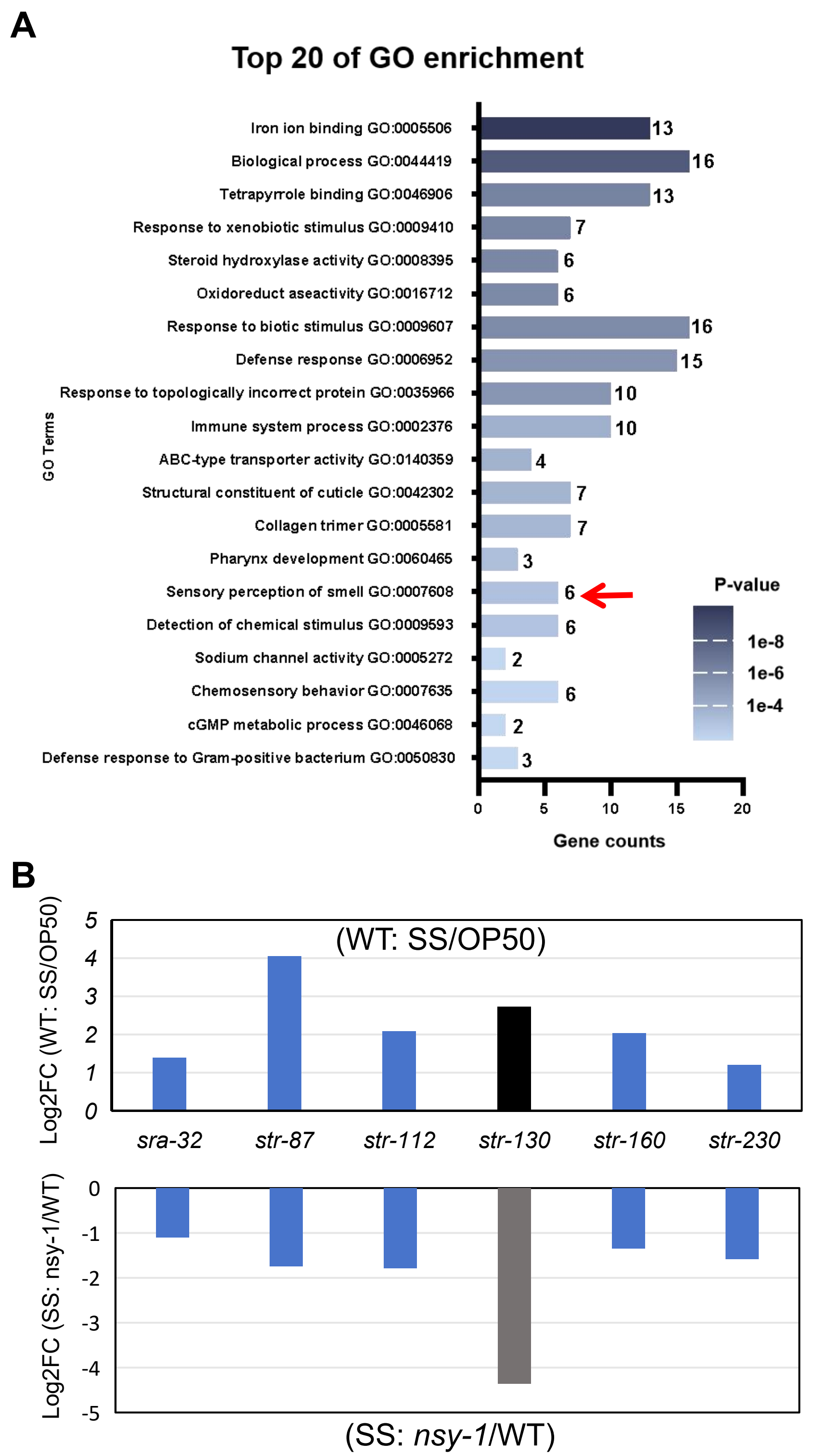


**Figure 4—figure supplement 1. *str-130* is induced in wild-type in response to SS, dependent on NSY-1.**

**(A)** GO enrichment analysis of 304 NSY-1-dependent candidate genes responding to SS (Supplementary file 2). Genes related to sensory perception (*sra-32, str-87, str-112, str-130, str-160, str-230*) are highlighted as enriched (red arrow).

**(B)** Relative mRNA expression levels of sensory perception-related genes (*sra-32, str-87, str-112, str-130, str-160, str-230*) extracted from RNA-seq data. These genes are induced in wild-type N2 animals in response to SS, but their expression is reduced in *nsy-1(ag3)* mutant animals under SS feeding conditions, indicating that the induction of these genes in wild-type in response to SS is dependent on NSY-1.

### Figure 4—figure supplement 2

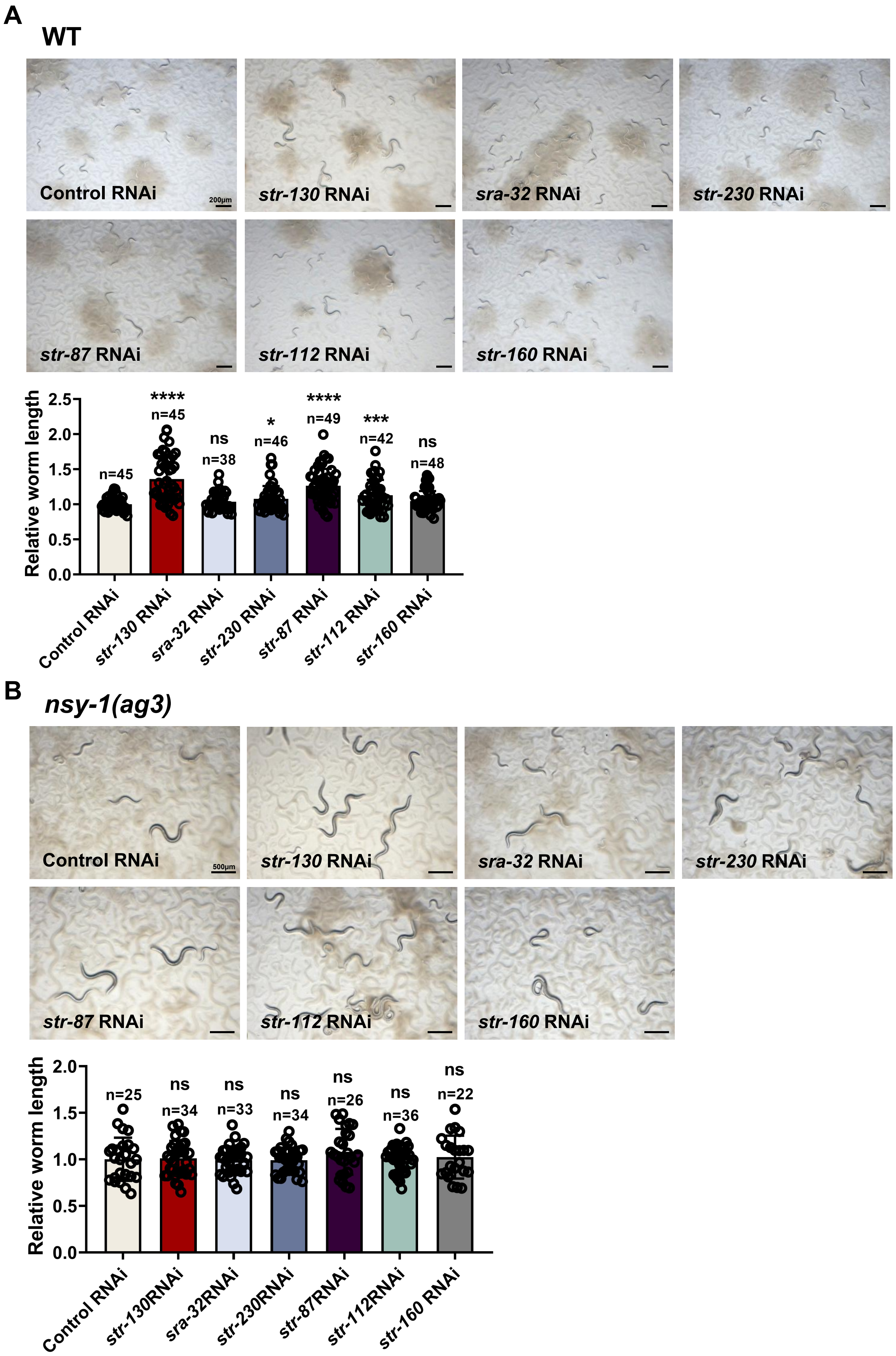

**Figure 4—figure supplement 2. *str-130* as the dominant effector of NSY-1-mediated SS response regulation.**

**(A)** Developmental progression of wild-type animals treated with control RNAi, *str-130* RNAi, *sra-32* RNAi, *str-230* RNAi, *str-87* RNAi, *str-112* RNAi or *str-160* RNAi grown on SS bacteria. Data are represented as mean ± SD. Scale bar = 200 μm. ****$p < 0.0001$; ***$p < 0.001$; *$p < 0.05$; n.s. not significant by Student's t-test (candidate RNAi vs Control RNAi). n= number of animals which were scored.

**(B)** Developmental progression of *nsy-1(ag3)* mutant animals treated with control RNAi, *str-130* RNAi, *sra-32* RNAi, *str-230* RNAi, *str-87* RNAi, *str-112* RNAi or str-160 RNAi grown on SS bacteria. Data are represented as mean ± SD. Scale bar= 500 μm. n.s. not significant by Student's t-test (candidate RNAi vs Control RNAi). n= number of animals which were scored.

### Figure 4—figure supplement 3

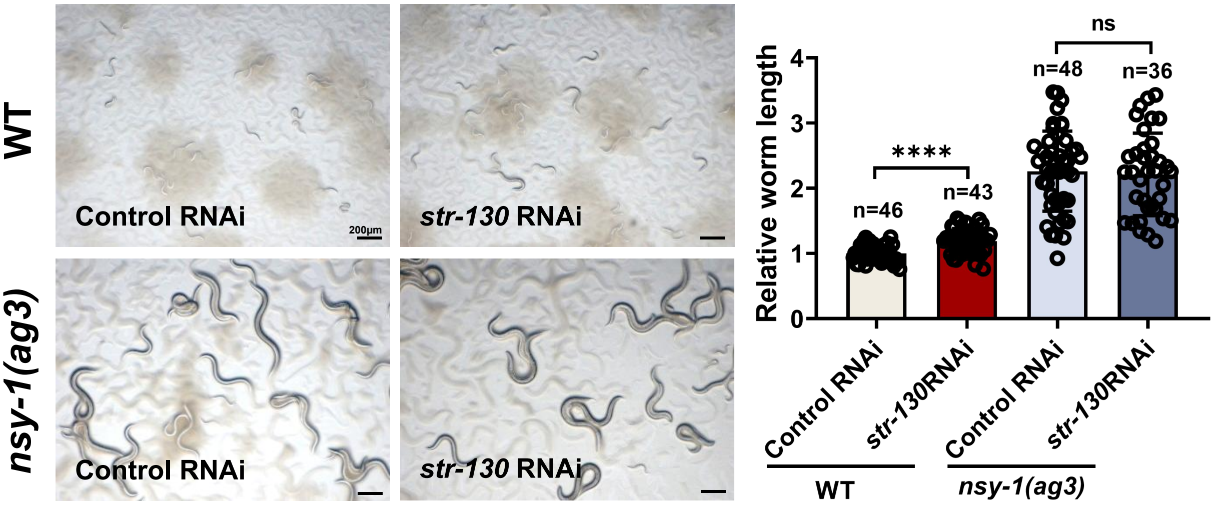


**Figure 4—figure supplement 3. NSY-1 inhibits SS digestion by inducing GPCR *str-130*.**

Developmental progression of wild-type or *nsy-1(ag3)* mutant animals treated with control RNAi or *str-130* RNAi grown on SS bacteria. Data are represented as mean ± SD. Scale bar = 200 μm. ****p < 0.0001; n.s. not significant by Student's t-test. n= number of animals which were scored.

### Figure 5—figure supplement 1

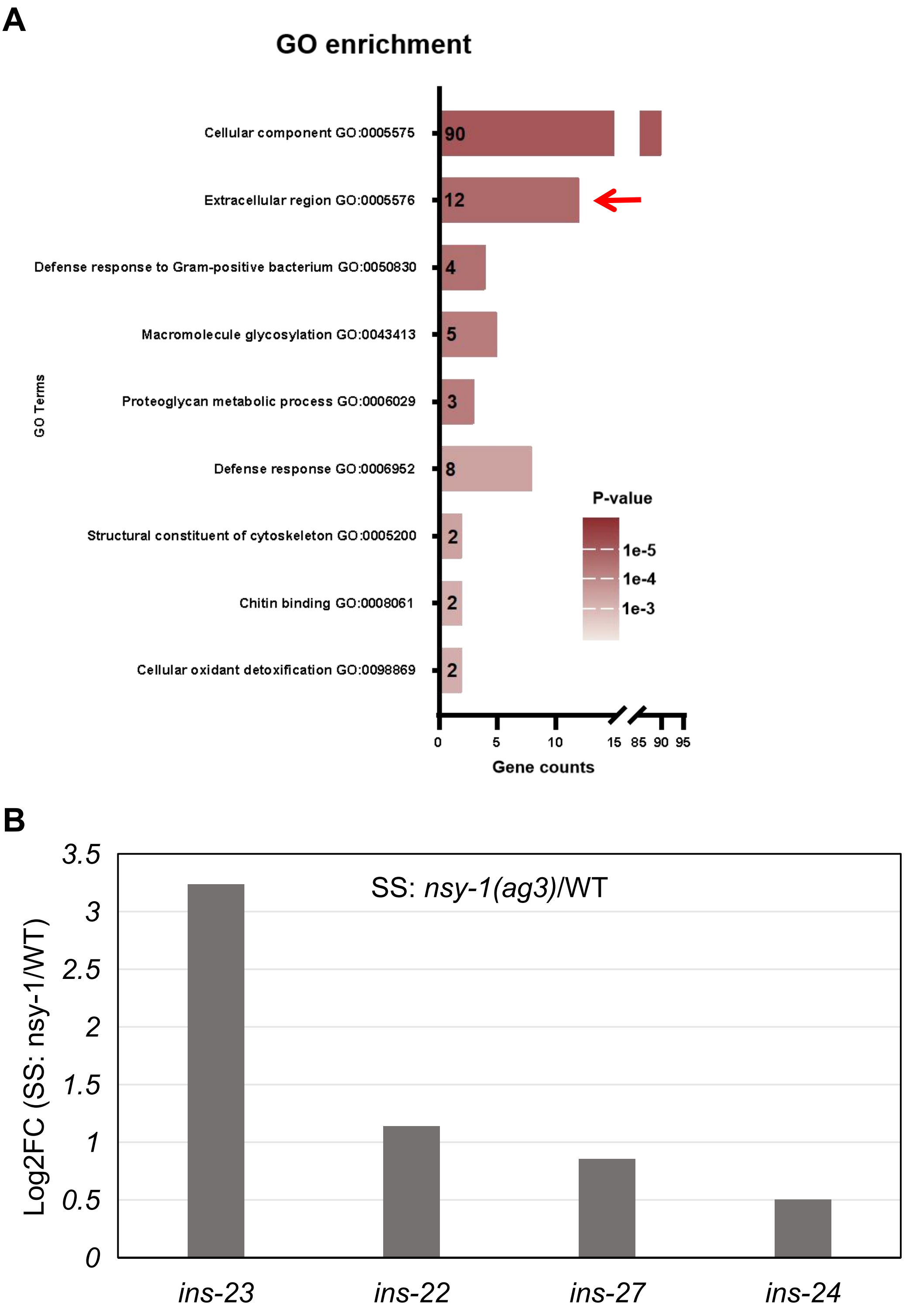


**Figure 5—figure supplement 1. *nsy-1* mutation induces the expression of insulin-related genes.**

**(A)** GO enrichment analysis of 308 genes induced by the nsy-1 mutation under SS feeding conditions (also see Supplementary file 2).

**(B)** Relative mRNA expression levels of insulin-related genes (*ins-23, ins-22, ins-27, ins-24*) extracted from RNA-seq data. These genes are upregulated in *nsy-1(ag3)* mutant animals under SS feeding conditions, which may contribute to SS digestion in the *nsy-1* mutant.

### Figure 5—figure supplement 2

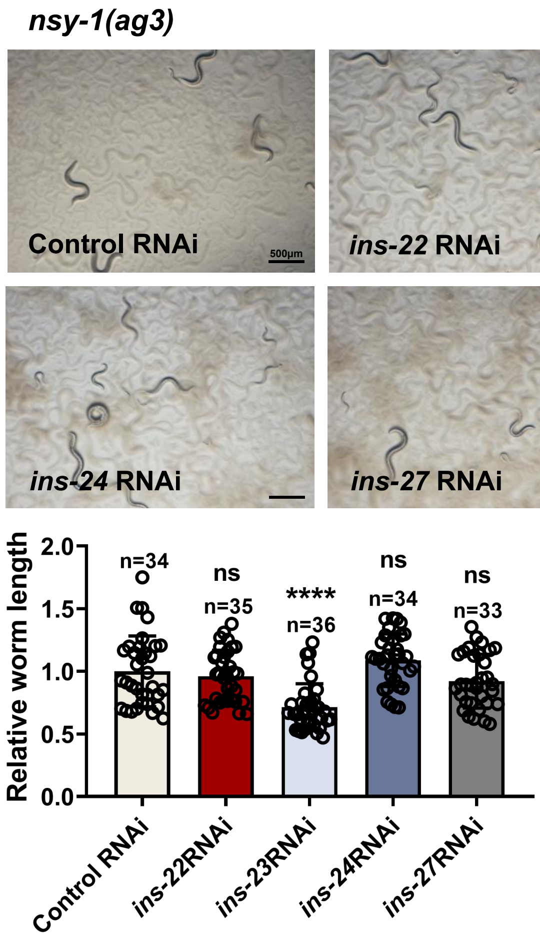


**Figure 5—figure supplement 2. NSY-1 mutation promotes SS digestion by inducing *ins-23*.** Developmental progression of *nsy-1(ag3)* mutant animals treated with control RNAi, *ins-22* RNAi; *ins-23* RNAi; *ins-24* RNAi; or *ins-27* RNAi grown on SS bacteria. Data are represented as mean ± SD. Scale bar = 500 µm. ****$p < 0.0001$; n.s. not significant by Student's t-test (candidate RNAi vs Control RNAi). n= number of animals which were scored.

### Figure 5—figure supplement 3

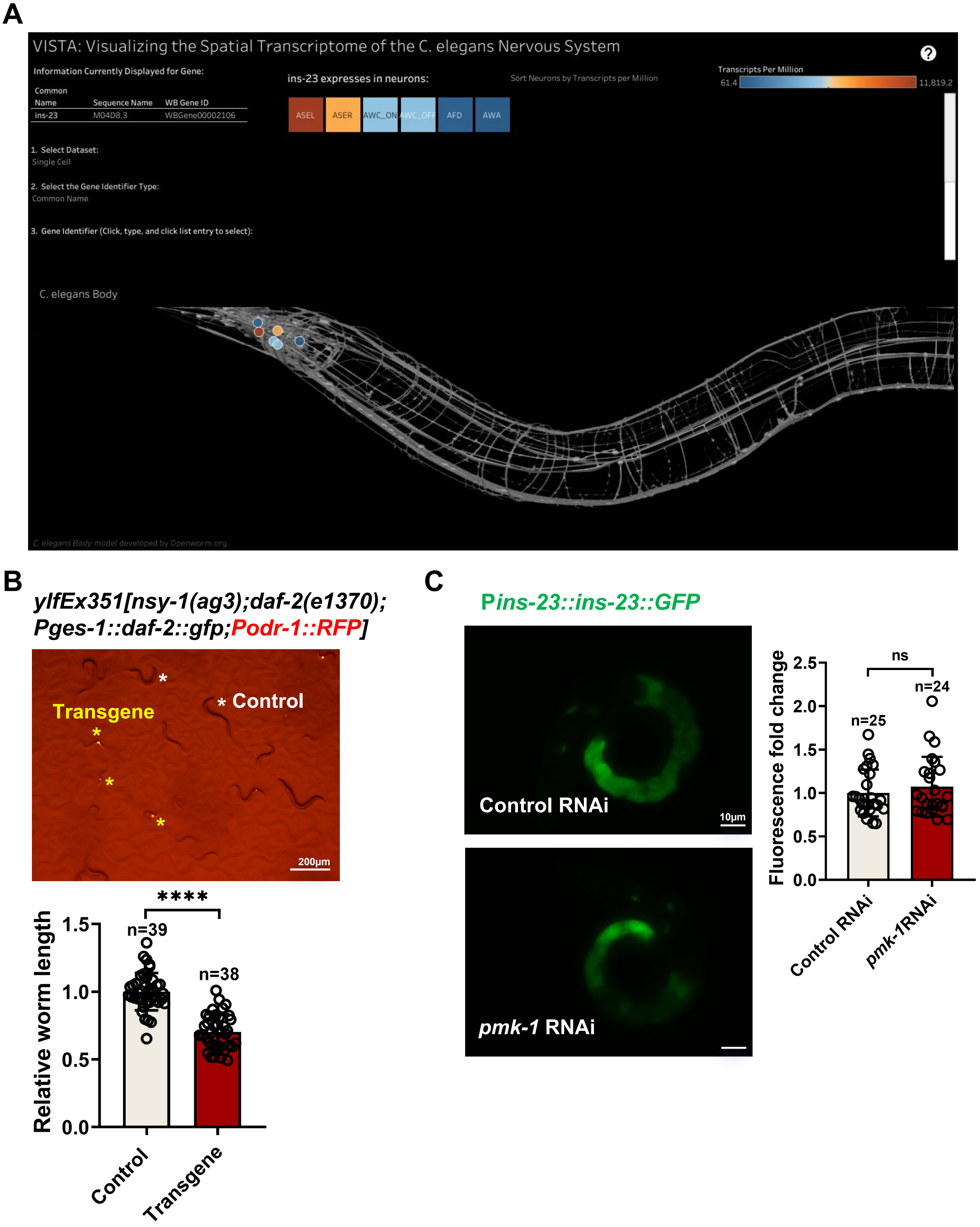

**Figure 5—figure supplement 3. INS-23 induction in *nsy-1* mutants promotes digestion independently of intestinal DAF-2 function.**

**(A)** CeNGEN predicts showing the head neurons expression pattern of *ins-23*. *ins-23* is expressed in AWC neurons.

**(B)** Developmental progression of *nsy-1(ag3);daf-2(e1370)* mutant worms carrying *Pges-1::daf-2::gfp* (intestine-specific expression) grown on SS bacteria. Control animals are labeled with white stars, and animals carrying the transgenes (rescued animals) are labeled with yellow stars. Data are represented as mean ± SD. Scale bar = 200 µm. ****$p < 0.0001$ by Student's t-test. n= number of animals which were scored.

**(C)** Microscopic images and quantitative data showing fluorescence of *Pins-23::ins-23::GFP* animals treated with control RNAi or *pmk-1* RNAi. Data are represented as mean ± SD. Scale bar = 10 µm. n.s. not significant by Student's t-test. n= number of animals which were scored.

#### Figure 6—figure supplement 1

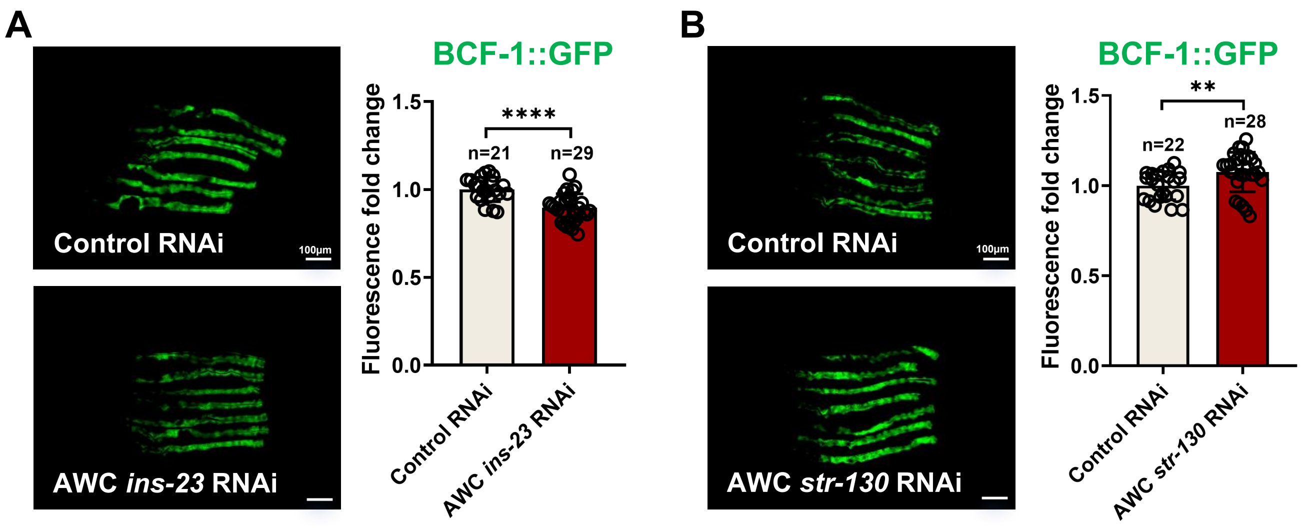


**Figure 6—figure supplement 1. INS-23 and STR-130 are functions cell-non-autonomously in AWC neurons to affect BCF-1 expression.**

**(A)** Microscopic images and quantitative data showing fluorescence of *Pbcf-1::bcf-1::GFP* animals treated with control RNAi or AWC *ins-23* RNAi. Data are represented as mean ± SD. Scale bar= 100 µm. ****$p < 0.0001$ by Student's t-test. n= number of animals which were scored.

**(B)** Microscopic images and quantitative data showing fluorescence of *Pbcf-1::bcf-1::GFP* animals treated with control RNAi or AWC *str-130* RNAi. Data are represented as mean ± SD. Scale bar= 100 µm. **$p < 0.01$ by Student's t-test. n= number of animals which were scored.
